# Winging it: Unveiling how hummingbirds alter their flying kinematics during molt

**DOI:** 10.1101/2023.11.21.568135

**Authors:** Andrés F. Díaz-Salazar, Felipe Garzón-Agudelo, Ashley Smiley, Carlos Daniel Cadena, Alejandro Rico-Guevara

## Abstract

Hummingbirds are well known for their hovering flight, one of the most energetically expensive modes of locomotion among animals. Molt is a costly event in the annual cycle, in which birds replace their feathers, including all their primary feathers which, in hummingbirds, comprise most of the area of the wing. Despite this, the effects of molt on hovering flight are not well known. Here, we examined high-speed videos (14 individuals of three species from the Colombian Andes recorded at 1200 FPS) comparing molting and non-molting hummingbirds’ wing kinematics and wingtip trajectories. We found that molting hummingbirds extended their wings in sharper angles during both downstroke and upstroke compared to non-molting individuals (10° vs 20°, and 15° vs 29°, respectively), while other flight parameters remained unchanged. Our findings show that hummingbirds are capable of sustaining hovering flight kinematics even under impressive wing area reductions by adjusting their wing flapping behavior.

## INTRODUCTION

Birds experience various developmentally demanding events throughout life, including reproduction, molt, and, in some cases, migration, which are especially limiting periods in the annual cycle (Bridge, 2011). Among these events, molt (i.e., the periodic shedding and replacement of feathers; Palmer, 1972) is relatively understudied in highly specialized flying birds like hummingbirds. Because birds exhibit increased metabolic rates during molting (Chai, 1997; Walsberg, 1983), molt periods are expected to be particularly challenging for hummingbirds, given their high mass-specific metabolic rates, which are the highest among vertebrates in association with their demanding hovering flight (Altshuler and Dudley, 2002; Suarez, 1992).

Furthermore, molt does not likely affect all avian groups in the same manner, given the high diversity of body plans among birds and their varied molt strategies (Chai and Dudley, 1999; Cherrel et al., 1994; Langston and Rohwer, 1996; Palmer, 1972; Swaddle and Witter, 1997). For instance, morphological changes during molt, such as wing gaps caused by feather shedding, are likely to affect the flight performance of birds. In the case of hummingbirds, unlike most birds, they produce lift during both the downstroke and the upstroke (Warrick et al., 2015), following an “insect-like” flying pattern. The main function of flight feathers is to work as airfoils that generate lift (Rayner, 1988). Therefore, a reduction in the airfoil area during molt is likely to decrease lift production, potentially resulting in energetic stress (Chai, 1997), and consequently affecting hovering kinematics (Achache et al., 2017; 2018).

A basic understanding of the effects of molt on hummingbird hovering kinematics is still lacking. Previous research showed no significant effects of molt on aerodynamic variables related to hovering aerodynamics, such as flapping frequency and stroke amplitude in the Ruby-throated Hummingbird (*Archilochus colubris;* Chai, 1997). However, the author cautioned that further analysis using high-speed video and vortex visualization techniques would be necessary to precisely detect any molt-related effects on flight performance (Chai, 1997). A recent study suggested that hummingbird wings may have reduced capacity to generate lift during the downstroke while moltingbased on vortex and force measurements compared to non-molting hummingbirds (Achache et al., 2018). However, these analyses were conducted on wing models rather than live, free-flying birds.

Hovering is an energetically expensive locomotion strategy (Sargent et al., 2021). Interestingly, hummingbirds can sustain their flight demands throughout their annual cycle despite losing up to 40% of total airfoil planform area during molt (Fig. S1). This begs the question of how they overcome the effects of molting to maintain hovering flight. A bird’s weight during flight is supported in proportion to its airfoil planform area. If weight remains static following a reduction in planform area (i.e., due to molting), there are some mechanical avenues that might allow individuals to compensate for lack of weight support through aerodynamic changes. First, we would expect to observe alterations in the flying pattern (i.e., the path followed by wings) of molting individuals compared to those with intact wing area. Second, compensating for area loss can be achieved by modifying body part angles or flapping frequency during flight (e.g., Fernandez et al., 2012). For example, one may predict molting hummingbirds to alter kinematic variables such as stroke amplitude, body positional angles, and flapping frequency. In this study, we aim to evaluate the effect of molt on the hovering kinematics and flying patterns of three species of hummingbirds in the Andes of Colombia, using high-speed video and geometric morphometric analysis. To our knowledge, this is the first empirical study on the flight kinematics of free-ranging hummingbirds during molt.

## MATERIALS AND METHODS

### Data collection

We captured high-speed videos of free-ranging hummingbirds at the Colibrí Gorriazul Research Station, located on the western slope of the eastern Cordillera de los Andes near Bogotá, Colombia at an elevation of 1700 m. Birds were captured and sampled using either mist nest or feeder traps. Prior to recording, we tagged hummingbirds with 7 mm PIT tags implanted subcutaneously between scapulae (Hou, Verdirame, and Welch, 2015) and then identified individuals with a Radio Frequency Identification (RFID) antenna. To allow for full wing motion during backward extension, the tag was placed between the thoracic and lumbar vertebrae, avoiding any obstruction (Bandivadekar et al., 2018).

Next, we used two FASTEC^®^ IL5 (Fastec Imaging, San Diego, CA, USA) high-speed cameras to record both molting and non-molting hummingbirds within a semi-controlled environment, i.e., a transparent Plexiglas chamber (90cm x 90 cm x 90 cm). The recessed chamber was connected to a laboratory window that opened to the outside, where feeders were strategically positioned to attract free-ranging hummingbirds. Inside the chamber, a 10 mL syringe was connected to an artificial flower, which provided a 20% sucrose solution *ad libitum*. Cameras were positioned 90° from each other, enabling simultaneous recording of both top and side views of the artificial flower and the visiting hummingbirds.

Hummingbirds voluntarily entered the chamber to access the artificial nectar provided through the syringe. Following an acclimation period (15 days approximately), we recorded high-speed videos at 1200 frames s^-1^ for individuals of three species (Indigo-capped Hummingbird *Saucerottia cyanifrons, n=5;* Rufous-tailed Hummingbird *Amazilia tzacatl, n=4;* and Black-throated Mango *Anthracothorax nigricollis, n=5*). We filmed 14 sets of videos (top and side view were taken as a single set), including five sets for molting birds and nine sets for non-molting birds. For our analysis, we exclusively focused on measuring static hovering flight between drinking bouts (excluding flying sequences where we detected horizontal movement of the individual). No measurements were taken while individuals were actively engaged in nectar consumption, rotating, or moving across different planes. We did not systematically assess particular stages of molt; instead, we simply recorded the presence or absence of wing molt. Furthermore, because our study emphasized free-flying individuals that went in and out from our study set, we were not able to measure their weight, and so we did not account for potential effects of body mass.

### Hovering Kinematics

To assess the impact of molt on hovering kinematics, we chose consistent body landmarks from individuals to look for alterations in their positional angles (Fig. 1A). First, each video was converted to a stack of images to conduct a frame-by-frame analysis. We analyzed five complete wingbeat cycles (i.e., one downstroke and one upstroke per cycle) for each set of synchronized videos. We used the software Fiji (ImageJ; Schindelin et al., 2012) to measure flapping frequency, stroke amplitude, and angles describing the position of selected body parts as indicators of flight kinematics (Fig. 1A, B) following previous studies (Achache et al., 2018; Altshuler et al., 2005; Chai, 1997). We measured the variables separately for each wingbeat cycle’s downstroke and upstroke.

**Figure 1.**
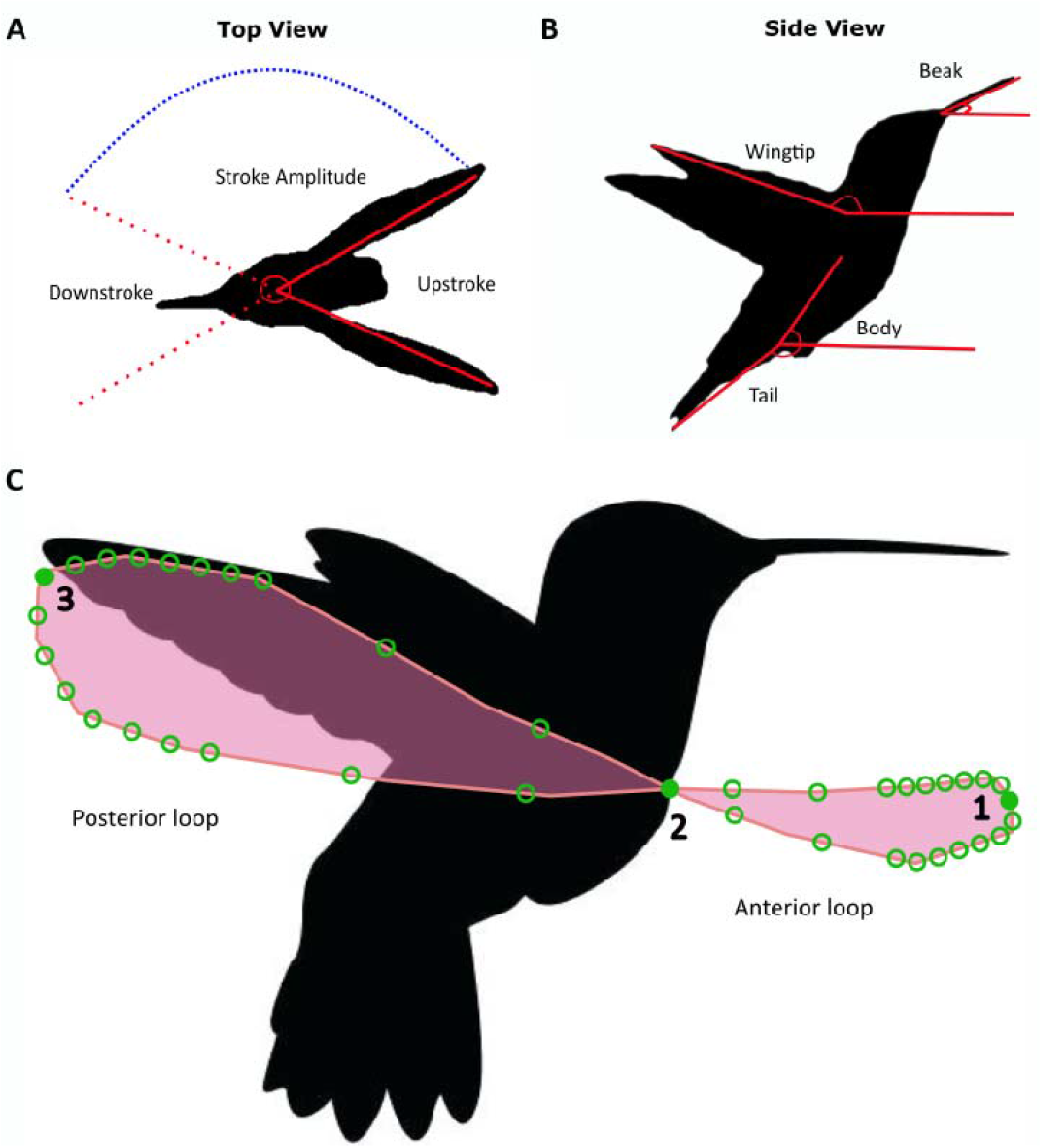
Kinematic variables located over the top view and side view. (A) Representation of the top view at maximum positional angle for the upstroke (solid red line) and maximum positional angle for the downstroke (red dotted line) (B) Body parts considered for the side view perspective. (C) Analemma (i.e. figure-eight pattern) formed by the wingtip trajectory during a complete wingbeat (pink line) and landmarks (numbered filled circles) and semilandmarks (open circles) that characterize it.

For the top-view kinematics, we quantified maximum and minimum wingtip positional angles (for downstroke and upstroke, respectively) as a measurement of wing opening and closing (Fig. 1A). For the side-view, we measured the positional angles of four body parts by digitizing points on the beak, tail, feet, and wingtips (Fig. 1B). Additionally, following Groom et al. (2018), we estimated flapping frequency by dividing the recording frame rate (1200 frames s^-1^) by the total number of frames that were necessary for a hummingbird to complete a single flap. The average was taken from five consecutive flap cycles on each video.

### Wingbeat patterns: Analemma size and shape

To assess whether molting hummingbirds exhibited altered hovering patterns compared to non-molting hummingbirds, we analyzed the lemniscate-shaped stroke cycle (figure-eight pattern) traced by wingtips during each wingbeat, hereafter referred to as an analemma (Fig. 1C). Using a two-dimensional geometric morphometric approach, we assessed differences in size and shape of analemmas between molting and non-molting hummingbirds from the side view video sets. For each video, we traced the wing tip trajectory over a minimum of three complete wingbeat cycles, using the plugin MtrackJ (Meijering et al., 2012) in the Fiji software (Schindelin et al., 2012). After digitizing the videos, we obtained a total of 85 analemmas, 30 for molting birds and 55 for non-molting birds.

To analyze the shape and size of the analemmas, we digitized 3 landmarks and 36 semilandmarks using TpsDig2 (Rohlf, 2021; Fig. 1C; see Supplementary Materials for details). We calculated the perimeter of each analemma by summing distances between landmarks and semilandmarks using the *interlmkdist* function in the *geomorph* package for R (Baken et al., 2021). Because the perimeter and centroid size were strongly correlated (r=0.99, *P*=0.0001), we focused solely on the perimeter as a size variable for subsequent analysis.

As a proxy for symmetry, we defined a length proportion corresponding to each analemma. We described its shape as consisting of two loops, one anterior and one posterior (Fig. 1C). We then calculated the ratio between the length of the anterior loop (distance between landmarks 1 and 2) and the overall length of the analemma (distance between landmarks 1 and 3; Fig. 1C). We obtained the shape variables (Procrustes tangent coordinates) through a Generalized Procrustes analysis (GPA; Rohlf, 1999) using the function *gpagen* from *geomorph*. This analysis effectively accounted for non-shape variation stemming from differences in position, scale, and orientation. For the GPA procedure, semilandmarks were allowed to slide under the criterion of bending energy minimization (Gunz and Mitteroecker, 2013; see Supplementary Materials for details). The resulting Procrustes coordinates were orthogonally projected from a curved space to a tangent space, yielding Procrustes tangent coordinates (Rohlf, 1999; Mitteroecker and Gunz, 2009; see Supplementary Materials for details).

Subsequently, we averaged perimeter and Procrustes tangent coordinates by individual. To obtain a representation of shape space, we performed a Principal Component Analysis (PCA) using the covariance matrix of the Procrustes tangent coordinates (*gm.prcomp* function from *geomorph*). We then generated wireframes to visually depict shape variation associated with the extremes of the PCA axes (Klingenberg, 2013).

### Statistical analyses

To statistically assess differences between molting and non-molting kinematics we conducted generalized linear models (GLMs) with the *glm* function from the stats package (R Core Team, 2012). We conducted separate GLM per variable and considered molt state and species as fixed effects to test for any influence on hovering kinematics.

We assessed the effect of molt on the analemma perimeter and the ratio of anterior loop to analemma length (symmetry proxy) with Mann-Whitney U tests. For analemma shape, we performed Procrustes ANOVA with permutation procedures (Goodall, 1991), where the Procrustes tangent coordinates (shape) were the response variable and molting state was the categorical variable (using the *procD.lm* function from *geomorph*). We also performed additional Procrustes ANOVA to assess allometry (shape∼perimeter) and the relation between shape and our proportional metric of symmetry (i.e., the ratio of anterior loop to analemma length). To quantify and compare the variation in analemma shape (Procrustes variance) between molting and non-molting birds (Zelditch et al., 2012), we used the *morphol.disparity* function from *geomorph*. All analyses were conducted in R version 4.2.1 (R Core Team, 2022; http://www.R-project.org/).

We assessed the influence of species on every kinematic and morphometric variable (results not shown). Although the variation was partly explained by interspecific differences, small sample sizes likely bias statistical results and thus we will not further explore the effect of species differences. We address this caveat of our work in the Discussion.

## RESULTS

### Hovering Kinematics

Based on the top-view video analysis, molting and non-molting hummingbirds differed in stroke amplitude, minimum positional angles (downstroke), and maximum positional angles (upstroke) (average change of 157° vs 138°, *P*<0.001; 10° vs 20°, *P*<0.001; and 15° vs 29°, *P*<0.004, respectively; Fig. 2, Table S1). Molting hummingbirds showed sharper terminal angles during their downstroke and upstroke, resulting in higher stroke amplitudes compared to non-molting hummingbirds (Fig. 2). In contrast, in the side view, we did not find any significant differences in positional angles between molting and non-molting birds (*P*>0.05 in all cases; Fig. S2, Table S2). Additionally, molting birds exhibited similar wing-flapping frequencies compared to non-molting birds (*P*=0.267; Fig. S3, Table S1).**Wingbeat patterns: Analemma size and shape** We observed no significant differences between molting and non-molting birds in the perimeter of the analemmas or in the ratio of the anterior loop to the length of the analemma (*P*=0.606, *P*=0.364, respectively; Fig. 3). Additionally, we did not find any significant allometric effect (R^2^ = 0.19, *P*=0.117).

**Figure 2.**
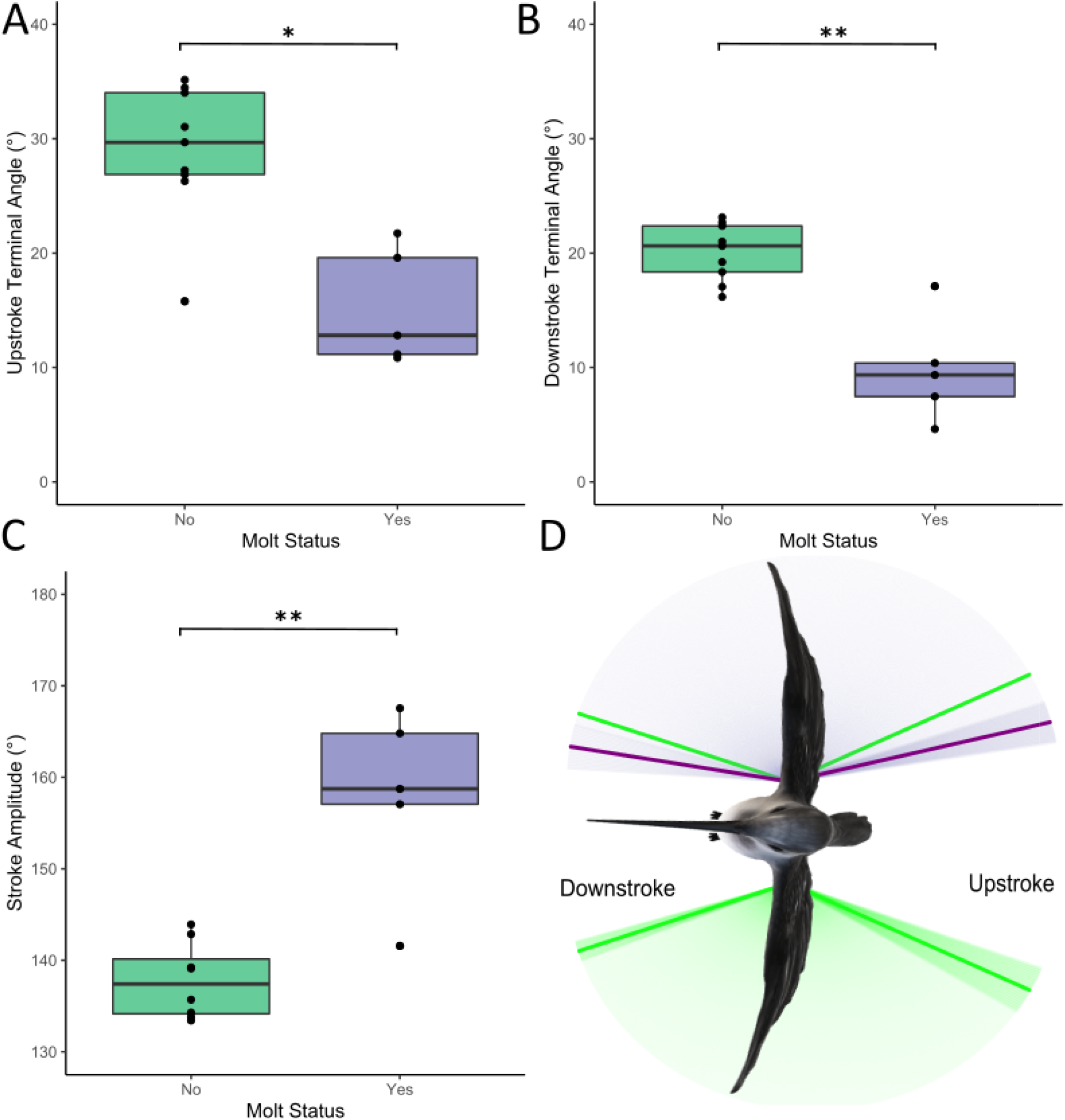
Top-view measurements revealing that molting hummingbirds showed higher stroke amplitudes. (A). Wing maximum positional angles (upstroke, B) and minimum positional angle (downstroke, C) were more acute in molting hummingbirds, suggesting hummingbirds got their wings closer in each wing beat during upstroke and downstroke during molt. As a result, (D) molting hummingbirds showed larger wingtip trajectories than non-molting hummingbirds (solid lines showing stroke angle average and shades represent their standard deviation). Purple represents molting and green non-molting.

**Figure 3.**
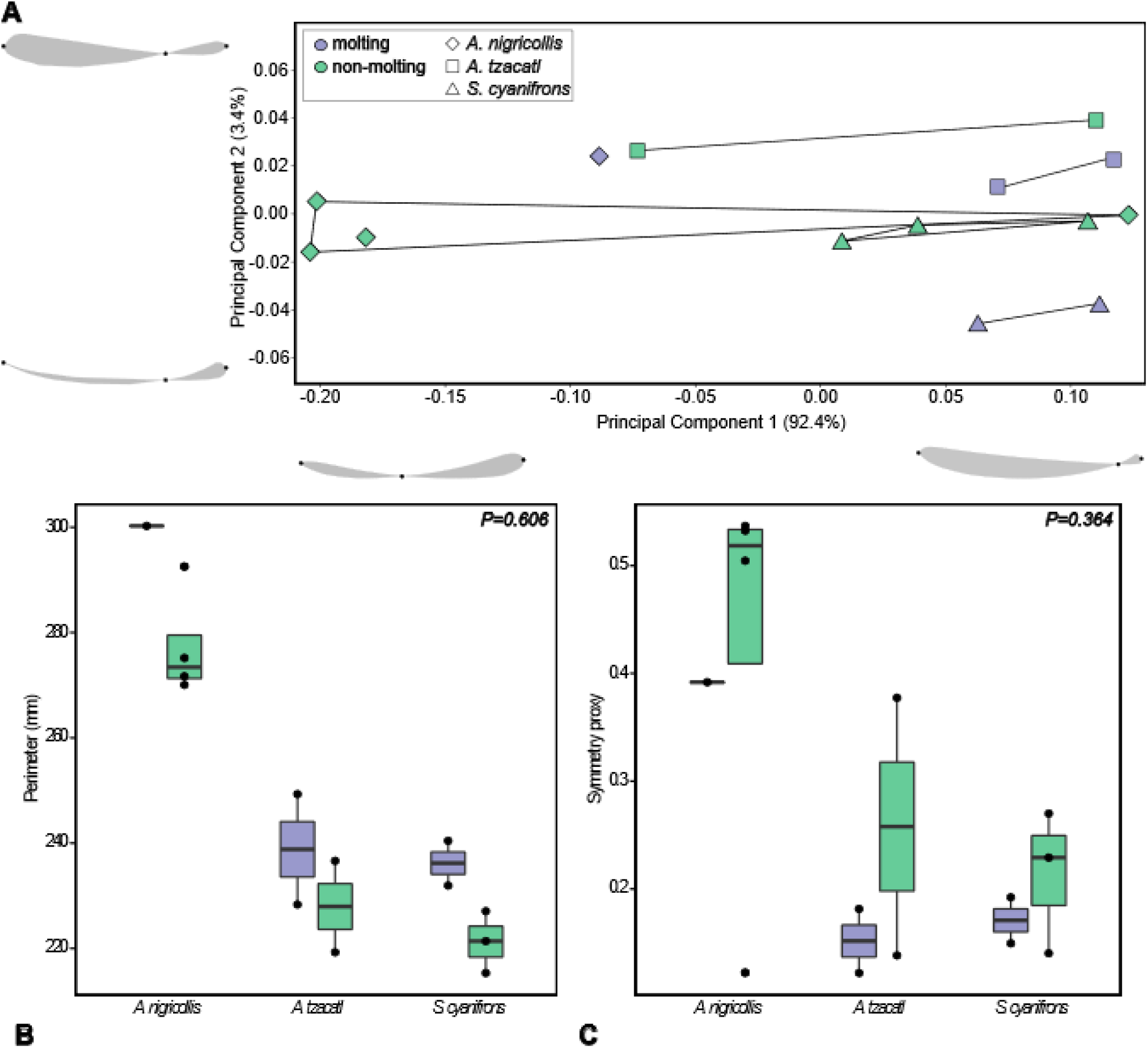
Molting status did not affect analemma size or shape. (A) First 2 principal components representing analemma shape space. Wireframes represent the shape variation associated with the extremes of the PCs. Semilandmarks are not shown. Wireframes have the same orientation as Fig. 1C. (B) Analemma perimeter and (C) Ratio of anterior loop to analemma length for molting and on-molting individuals per species. Statistical differences were evaluated between all molting and non-molting individuals (P values in B and C), intraspecific differences were not tested. Purple represents molting and green non-molting individuals.

The principal component analysis (PCA) revealed that the first two principal components combined accounted for 95.83% of the total variation in the shape of the analemmas (PC1=92.44%, PC2=3.39%; Fig. 3). The Procrustes ANOVA indicated that the symmetry proxy explained the majority of analemma shape variation (R^2^ = 0.92, *P*=0.001), representing a simple alternative to characterize analemma shape. PC1 showed a strong association with this metric. Specifically, analemmas were more symmetric toward the negative side of PC1, corresponding to higher ratio values, and had relatively shorter anterior loops toward the positive side of PC1, corresponding to lower ratio values. PC2 described the proximity of the downstroke and upstroke trajectories, particularly in the posterior loop of the analemmas. There was no clear separation in the analemma shape space between molting and non-molting hummingbirds, as indicated by the Procrustes ANOVA (R^2^ = 0.11, *P*=0.203; Fig. 3).

Hummingbirds with intact wing area exhibited more variance in analemma shape than molting birds, but the difference in variances was not significant (Procrustes variance was 0.0176 in non-molting birds and 0.0073 in molting birds, *P*=0.118). The same pattern of variation can be visualized per species along PC1 and in the boxplot of the ratio of the anterior loop to the analemma length (Fig. 3).

## DISCUSSION

We examined how molt affects the hovering kinematics and the wingbeat pattern of free-ranging hummingbirds. Our findings suggest that hummingbirds may overcome flight limitations imposed by molt through adjustments in wing angles during both the upstroke and downstroke. Hummingbird wings trace an analemma-like shape during wingbeats, but remarkably, these trajectories remained similar in molting and non-molting hummingbirds.

We found that molting hummingbirds exhibit greater angles of amplitude on both the downstroke and upstroke than non-molting hummingbirds when seen from above (Fig. 2). This means that molting individuals carry their wings a longer distance to complete a single flap than non-molting individuals in both downstroke and upstroke. Wider stroke amplitudes result in a greater production of aerodynamic forces, leading to increased lift (Altshuler and Dudley, 2002; Skandalis et al., 2017). Because feather shedding generates gaps in the wings, which results in a distortion of the optimal wing shape for lift generation (Achache et al., 2018), molting hummingbirds appear to offset these morphological alterations by extending their wings farther in each stroke. Because wing-area reduction leads to an increase in wing-loading (i.e., the relationship between wing area and body mass), higher stroke amplitudes may compensate for the loss of lift (Chai and Millard, 1997), thus alleviating the morphological limitations imposed by molt.

Conversely, apart from the changes in stroke amplitude observed from a top view, molting hummingbirds did not alter the other kinematic and positional variables relative to non-molting hummingbirds when observed from the side. Despite experiencing substantial reductions in airfoil area, molting individuals managed to maintain consistent positional angles of their wings, bill, body, and tail. Additionally, during hovering, molting hummingbirds maintained their wingtip trajectory (wingbeat pattern), as shown in our geometric morphometric analysis. We expected to observe alterations in the eight-shaped trajectory (analemma) traced by hummingbirds’ wingtips between molting and non-molting hummingbirds, which would reflect a compensatory mechanism against molt. However, we found no differences between molting and non-molting birds in the shape or perimeter of their analemmas. Nevertheless, for every species, molting birds seem to exhibit less shape variation compared to non-molting birds (Fig. 3A, C; Fig. S4). This pattern warrants further exploration, as it may suggest that molt imposes restrictions on the variation of the analemma shape, potentially influencing aerodynamic performance. Furthermore, although the molting state did not have a significant effect on the perimeter of the analemma, in each species molting birds consistently exhibited higher maximum perimeter values compared to non-molting birds (Fig. 3B). This finding aligns with the increased stroke amplitudes observed in the kinematic analysis.

Wing area reductions would normally result in greater wing loading, which could be expected to lead to locomotor compensations (Swaddle and Witter, 1997). One possible behavioral compensatory mechanism could be to reduce their body mass to alleviate kinematic constraints (Achache et al., 2018; Chai, 1997). However, it remains unclear whether mass reduction occurs across hummingbird species and sexes, and to what extent body mass modulation can be achieved in the wild. Furthermore, it is uncertain whether hummingbirds can voluntarily control their body mass to compensate for wing area reduction while maintaining their ability to generate lift, or if changes in mass instead reflect increased physiological demands of feather synthesis (Swaddle and Witter, 1997). Because we did not measure body mass, we are unable to determine whether the studied species reduce their mass during molt.

Another way to compensate for potentially detrimental effects of molt on hovering may be to increase wing flapping frequency, thereby reaching equivalent aerodynamic forces to those produced via high-amplitude flapping (Altshuler et al., 2005; Fernández et al., 2012; Rajabi et al., 2020). However, we found no differences in flapping frequency between molting and non-molting hummingbirds. In contrast, molting birds traced larger wing trajectories maintaining on average the same flapping frequencies compared to non-molting birds, which would imply higher flapping velocities to achieve farther wing trajectories.

Our study, constrained by the available sample sizes, did not extensively explore interspecific nor intraspecific variation of strategies to cope with aerodynamic restrictions related to molt (Wilcox and Clark, 2022). Hummingbirds, characterized by diverse ecological strategies and evolutionary pathways, exhibit significant variation in wing morphology and flight physiology (Altshuler and Dudley, 2002). Consequently, compensatory mechanisms reducing the impact of molt may vary among different clades. Recognizing the importance of robust group estimates, including means and variances, to reach meaningful conclusions (Cardini et al., 2015), we encourage the use of larger sample sizes to perform interspecific analyses. Additionally, we acknowledge that our geometric morphometric analysis may have underestimated the size and shape variation of analemmas by disregarding the third dimension (Cardini, 2014). Furthermore, our analysis did not assess the individual effects of single feather gaps. During molt, birds are likely to experience different effects on flying performance and energetic expenditure depending on which feathers are shedding or being replaced. Although there is a lack of information regarding how molt influences the life history of hummingbirds, an interesting fact is that the molt pattern is reversed for primaries IX and X in hummingbirds, possibly as an adaptation to mitigate the effects of molt on lift performance (Baltosser, 1995; Chai, 1997; Stiles, 1995). We note that our analysis was limited to hovering flight and did not consider potential implications on forward flight, acceleration, and maneuverability, nevertheless; it shows how these highly adapted-to-flight birds overcome molt’s morphological changes. Furthermore, we encourage future studies to delve into a better understanding of the effects of molt on hummingbird physiology, behavior, and ecology.

## Acknowledgements

We thank the team of researchers at Colibrí Gorriazul Research Center for their help with the methodology set up during the field season, the members of the Evolvert Lab from the Universidad de los Andes and the Behavioral Ecophysics Lab from the University of Washington for their insights on the manuscript. To Nelson Falcón for his suggestions on the geometric morphometrics analysis, and to Michael Dillon for sharing his R Script for the stroke amplitude figure. And finally, to Lucero Simbaqueba, Parmenio Simbaqueba, and Mary Simbaqueba for their help in the maintenance of feeders and work at Colibrí Gorriazul Research Center.

## Author contributions

Conceptualization: A.D.-S., A.R.-G., C.D.C.; Methodology: A.D.-S., A.R.-G., F.G.-A; Formal Analysis: A.D.-S., A.R.-G., F.G.-A; Investigation: A.D.-S., A.R.-G., F.G.-A., A.S.; Writing original draft: A.D.-S., A.R.-G., C.D.C.; Writing –review & editing: A.D.-S., A.R.-G., F.G.-A., C.D.C., A.S.; Visualization: A.D.-S., F.G-A.; Supervision: A.R.-G., C.D.C.

## Ethics

Hummingbirds tagging procedure was approved by Institutional Committee for the Use and Care of Laboratory Animals (CICUAL) from the Universidad de los Andes: C.FUA_19-002.

## 11. REFERENCES

Achache, Y., Sapir, N., & Elimelech, Y. (2017). Hovering hummingbird wing aerodynamics during the annual cycle. I. complete wing. Royal Society Open Science, 4(8), 1–8. 10.1098/rsos.170183

Achache, Y., Sapir, N., & Elimelech, Y. (2018). Hovering hummingbird wing aerodynamics during the annual cycle. II. Implications of wing feather molt. Royal Society Open Science, 5(2), 1–8. 10.1098/rsos.171766

Altshuler, D. L., Dickson, W. B., Vance, J. T., Roberts, S. P., & Dickinson, M. H. (2005). Short-amplitude high-frequency wing strokes determine the aerodynamics of honeybee flight. Proceedings of the National Academy of Sciences, 102(50), 18213–18218. 10.1073/pnas.0506590102

Altshuler, D. L., Dudley, R., Heredia, S. M., & McGuire, J. A. (2010). Allometry of hummingbird lifting performance. Journal of Experimental Biology, 213(5), 725–734. 10.1242/jeb.037002

Altshuler, D. L., & Dudley, R. (2002). The ecological and evolutionary interface of hummingbird flight physiology. Journal of Experimental Biology, 205, 2325–2336.

Baken, E. K., Collyer, M. L., Kaliontzopoulou, A., & Adams, D. C. (2021). geomorph v4.0 and gmShiny: Enhanced analytics and a new graphical interface for a comprehensive morphometric experience. Methods in Ecology and Evolution, 12(12), 2355–2363. 10.1111/2041-210X.13723

Baltosser, W. H. (1995). Annual molt in Ruby-throated and Black-chinned hummingbirds. The Condor, 97(2), 484–491. 10.2307/1369034

Bandivadekar, R. R., Pandit, P. S., Sollmann, R., Thomas, M. J., Logan, S. M., Brown, J. C., … Tell, L. A. (2018). Use of RFID technology to characterize feeder visitations and contact network of hummingbirds in urban habitats. PLoS ONE, 13(12), 1–25. 10.1371/journal.pone.0208057

Bridge, E. S. (2011). Mind the Gaps: What’s Missing of Feather Molt, 113(1), 1–4. 10.1525/cond.2011.100228

Cardini, A. (2014). Missing the third dimension in geometric morphometrics: how to assess if 2D images really are a good proxy for 3D structures? Hystrix, 25, 73–81. 10.4404/hystrix-25.2-10993

Cardini, A., Seetah, K., & Barker, G. (2015). How many specimens do I need? Sampling error in geometric morphometrics: testing the sensitivity of means and variances in simple randomized selection experiments. Zoomorphology, 134, 149–163. 10.1007/s00435-015-0253-z

Chai, P., & Millard, D. (1997). Flight and size constraints: hovering performance of large hummingbirds under maximal loading. The Journal of Experimental Biology, 200(Pt 21), 2757–2763.

Chai, Peng. (1997). Hummingbird hovering energetics during moult of primary flight feathers. Journal of Experimental Biology, 200(10), 1527–1536.

Chai, Peng, Altshuler, D. L., Stephens, D. B., & Dillon, M. E. (1999). Maximal Horizontal Flight Performance of Hummingbirds: Effects of Body Mass and Molt. Physiological and Biochemical Zoology, 72(2), 145–155.

Chai, Peng, & Dudley, R. (1999). Maximum Flight Performance of Hummingbirds: Capacities, Constraints, and Trade-Offs. The American Naturalist, 153(4), 398–411. 10.1086/303179

Feinsinger, P., & Chaplin, S. B. (1975). On the relationship between wing disc loading and foraging strategy in hummingbirds. The American Naturalist, 109(966), 217–224.

Goodall, C. (1991). Procrustes methods in the statistical analysis of shape. Journal of the Royal Statistical Society. Series B (Methodological), 53, 285–339. 10.1111/j.2517-6161.1991.tb01825.x

Greenewalt, C. (1960). The wings of insects and birds as mechanical oscillators. Proceedings of the American Philosophical Society, 605–611. Retrieved from https://www.jstor.org/stable/985536

Groom, D. J. E., Toledo, M. C. B., Powers, D. R., Tobalske, B. W., & Welch, K. C. (2018). Integrating morphology and kinematics in the scaling of hummingbird hovering metabolic rate and efficiency. Proceedings. Biological Sciences, 285(1873), 20172011. 10.1098/rspb.2017.2011

Gunz, P., & Mitteroecker, P. (2013). Semilandmarks: a method for quantifying curves and surfaces. Hystrix, 24(1), 103–109. 10.4404/hystrix-24.1-6292

Hedrick, T. L., Tobalske, B. W., Ros, I. G., Warrick, D. R., & Biewener, A. A. (2012). Morphological and kinematic basis of the hummingbird flight stroke: scaling of flight muscle transmission ratio. Proceedings of the Royal Society B: Biological Sciences, 279(1735), 1986– 1992. 10.1098/rspb.2011.2238

Hou, L., Verdirame, M., & Welch, K. C. (2015). Automated tracking of wild hummingbird mass and energetics over multiple time scales using radio frequency identification (RFID) technology. Journal of Avian Biology, 46(1), 1–8. 10.1111/jav.00478

Klingenberg, C. P. (2013). Visualizations in geometric morphometrics: how to read and how to make graphs showing shape changes. Hystrix, 24(1), 15. 10.4404/hystrix-24.1-7691

Meijering, E., Dzyubachyk, O., & Smal, I. (2012). Methods for cell and particle tracking. Methods in Enzymology, 504, 183–200. 10.1016/B978-0-12-391857-4.00009-4

Mitteroecker, P., & Gunz, P. (2009). Advances in geometric morphometrics. Evolutionary Biology, 36, 235–247. 10.1007/s11692-009-9055-x

Palmer, R. S. (1972). Patterns of molting. Avian Biology, 2, 65–101.

Rayner, J. M. (1988). Form and function in avian flight. In Current Ornithology (pp. 1-66).

Boston, MA: Springer US. R Core Team (2022). R: A language and environment for statistical computing. R Foundation for Statistical Computing, Vienna, Austria. https://www.R-project.org/

Rohlf, F. J. (1999). Shape statistics: Procrustes superimpositions and tangent spaces. Journal of Classification, 16, 197–223. 10.1007/s003579900054

Rohlf, F. J. (2021). tpsDig, digitize landmarks and outlines, version 2.32. Department of Ecology and Evolution, State University of New York at Stony Brook.

Schindelin, J., Arganda-Carreras, I., Frise, E., Kaynig, V., Longair, M., Pietzsch, T., Preibisch, S., Rueden, C., Saalfeld, S., Schmid, B., et al. (2012). Fiji: an open-source platform for biological-image analysis. Nature Methods, 9(7), 676-682. 10.1038/nmeth.2019

Skandalis, D. A., Segre, P. S., Bahlman, J. W., Groom, D. J. E., Welch, K. C., Witt, C. C., & Altshuler, D. L. (2017). The biomechanical origin of extreme wing allometry in hummingbirds. Nature Communications, 8(1), 1–9.

Stiles, F. G. (1995). Intraspecific and interspecific variation in molt patterns of some tropical hummingbirds. The Auk, 112(1), 118. 10.2307/4088772

Suarez, R. K. (1992). Hummingbird flight: Sustaining the highest mass-specific metabolic rates among vertebrates. Experientia, 48(6), 565–570. 10.1007/BF01920240

Swaddle, J. P., & Witter, M. S. (1997). The effects of molt on the flight performance, body mass, and behavior of European starlings (Sturnus vulgaris): an experimental approach. Canadian Journal of Zoology, 75(7), 1135–1146. 10.1139/z97-136

Tobalske, B. W., Warrick, D. R., Clark, C. J., Powers, D. R., Hedrick, T. L., Hyder, G. A., & Biewener, A. A. (2007). Three-dimensional kinematics of hummingbird flight. Journal of Experimental Biology, 210(13), 2368–2382. 10.1242/jeb.005686

Walsberg, G. (1983). Avian ecological energetics. In “Avian Biology VII.” DS Farner, JR King, and KC Parks, Eds., 7.

Warrick, D. R., Tobalske, B. W., & Powers, D. R. (2005). Aerodynamics of the Hovering Hummingbird. 10.1038/nature03647

Weis-Fogh, T. (1972). Energetics of hovering flight in hummingbirds and in Drosophila. Journal of Experimental Biology, 56, 79–104.

Zelditch, M. L., Swiderski, D. L., & Sheets, H. D. (2012). Geometric Morphometrics for Biologists: A Primer. Academic Press.

